# Membrane extraction in native lipid nanodiscs reveals dynamic regulation of Cdc42 complexes during cell polarization

**DOI:** 10.1101/2023.06.12.544590

**Authors:** Lars N. Deutz, Sena Sarıkaya, Daniel J. Dickinson

**Affiliations:** Department of Molecular Biosciences The University of Texas at Austin 2415 Speedway, PAT 206 Austin, TX 78712

## Abstract

Embryonic development requires the establishment of cell polarity to enable cell fate segregation and tissue morphogenesis. This process is regulated by Par complex proteins, which partition into polarized membrane domains and direct downstream polarized cell behaviors. The kinase aPKC (along with its cofactor Par6) is a key member of this network and can be recruited to the plasma membrane by either the small GTPase Cdc42 or the scaffolding protein Par3. Although *in vitro* interactions among these proteins are well established, much is still unknown about the complexes they form during development. Here, to enable the study of membrane-associated complexes *in vivo,* we used a maleic acid copolymer to rapidly isolate membrane proteins from single *C. elegans* zygotes into lipid nanodiscs. We show that native lipid nanodisc formation enables detection of endogenous complexes involving Cdc42, which are undetectable when cells are lysed in detergent. We found that Cdc42 interacts more strongly with aPKC/Par6 during polarity maintenance than polarity establishment, two developmental stages that are separated by only a few minutes. We further show that Cdc42 and Par3 do not bind aPKC/Par6 simultaneously, confirming recent *in vitro* findings in an *in vivo* context. Our findings establish a new tool for studying membrane-associated signaling complexes and reveal an unexpected mode of polarity regulation via Cdc42.

## Introduction

Embryonic development requires the effective partitioning of cellular components to separate poles of the cell, a phenomenon referred to as cell polarity. Proteins in the Partitioning defective (Par) system have been identified to regulate this behavior (Goldstein and Macara, 2007; Lang and Munro, 2017). Atypical Protein Kinase C (aPKC) is an essential, evolutionarily conserved component of the Par system that phosphorylates cytoskeletal proteins, cell fate determinants, and other targets that execute polarized cell behaviors. aPKC forms a tight complex with its cofactor Par6 (Joberty et al., 2000; Lin et al., 2000; Qiu et al., 2000; Sarıkaya and Dickinson, 2021). The aPKC/Par6 heterodimer can be recruited to the plasma membrane, where it is thought to act, by the scaffolding protein Par3 and/or the small GTPase Cdc42 (Izumi et al., 1998; Wodarz et al., 2000; Lin et al., 2000; Qiu et al., 2000; Joberty et al., 2000; Johansson et al., 2000; Petronczki and Knoblich, 2000; Beers and Kemphues, 2006).

The *C. elegans* zygote is a simple and well-studied model of animal cell polarity. In this system, polarization occurs via distinct establishment and maintenance phases (Cuenca et al., 2003). Polarity establishment occurs via actomyosin-driven cortical flows on the plasma membrane, which transport Par3 oligomers along with bound Par complexes (aPKC/Par6) to establish the anterior domain of the cell (Munro et al., 2004; Goehring et al., 2011; Sailer et al., 2015; Dickinson et al., 2017; Chang and Dickinson, 2022; Illukkumbura et al., 2023). During polarity maintenance, Par3 clusters disassociate, and cortical levels of Par3 are reduced, but aPKC/Par6 complexes remain at the anterior cortical domain (Dickinson et al., 2017; Rodriguez et al., 2017; Wang et al., 2017). Anteriorly-localized aPKC plays an essential role in maintaining polarity by antagonizing the posterior Par proteins PAR-1 and PAR-2 (Watts et al., 1996; Tabuse et al., 1998; Hao et al., 2006; Motegi et al., 2011; Folkmann and Seydoux, 2019). Cdc42 is genetically implicated in polarity maintenance and is a likely candidate to stabilize active aPKC at the anterior plasma membrane (Gotta et al., 2001; Kay and Hunter, 2001; Motegi and Sugimoto, 2006; Rodriguez et al., 2017).

The exact function of Cdc42 in polarity regulation has been most thoroughly studied in yeasts (Pichaud et al., 2019). In these organisms, GTP-bound Cdc42 accumulates at the nascent bud site (in *S. cerevisiae*) or the growing cell poles (in *S. pombe*) through a positive feedback cycle in which cortical Cdc42 recruits guanine exchange factors (GEF) to further activate Cdc42 (Woods and Lew, 2019). Localized activation of Cdc42, via GTP loading, is thought to be the key event leading to cell polarization (Miller et al., 2020).

Activation of Cdc42 by GEFs is also thought to be critical for its function in animal cell polarity, including in *C. elegans (Kumfer et al., 2010)*. Par6 binds directly to Cdc42 in a GTP-dependent manner, via a CRIB-PDZ domain that comprises the C-terminal half of Par6 (Qiu et al., 2000; Garrard et al., 2003). The full-length aPKC/Par6 heterodimer can also bind to Cdc42 (Vargas and Prehoda, 2023). *In vitro,* Cdc42 and Par3 appear to compete for binding to aPKC/Par6, suggesting that aPKC/Par6 may be recruited to the plasma membrane via two distinct mechanisms, one involving Par3 and one involving active Cdc42 (Vargas and Prehoda, 2023). A model has emerged in the literature in which aPKC/Par6 complexes are bound and transported by Par3 during polarity establishment, then “handed off” to Cdc42 during polarity maintenance (Rodriguez et al., 2017). Though this model is appealing, it has not yet been tested via direct biochemical experiments. Alternatively, the existence of a quaternary complex containing Par3, Par6, aPKC, and Cdc42 has been reported in cells overexpressing a GTP-locked, constitutively active mutant of Cdc42 (Joberty et al., 2000), but evidence for a quaternary complex involving native proteins *in vivo* is lacking.

To better understand how Par polarity complexes and other signaling complexes are assembled and regulated during development, it is essential to study these complexes in a state that is as close as possible to their native *in vivo* environment. With this goal in mind, our lab has developed a quantitative biochemical approach, termed single-cell, single-molecule pull-down (sc-SiMPull) that can measure the abundance of protein complexes in single, precisely staged *C. elegans* zygotes (Dickinson et al., 2017; Sarıkaya and Dickinson, 2021). In brief, sc-SiMPull entails rapidly lysing a cell or embryo, capturing endogenously tagged protein complexes on the surface of a coverslip, and visualizing these complexes using single-molecule TIRF microscopy. We have previously shown that this method can reveal the abundance, stoichiometry, stability, and developmental regulation of native protein complexes (Dickinson et al., 2017; Sarıkaya and Dickinson, 2021).

In the present study, we employ sc-SiMPull to study the interaction between Cdc42 and Par6 in single-cell *C. elegans* embryos and reveal how Cdc42/aPKC/Par6 binding is differentially regulated at different stages of polarity. We found that detergent, which we have typically included in our sc-SiMPull experiments to solubilize membranes, disrupts native Cdc42/Par6 interactions; thus, these proteins require an intact lipid membrane in order to interact *in vivo*. To enable extraction and quantification of intact Cdc42/Par6 complexes, we adopted an alternative membrane solubilizing agent: a lipid nanodisc-forming maleic-acid copolymer that has been shown to stabilize native proteins in a lipid bilayer environment (Oluwole et al., 2017a; b). We show that these polymers allow rapid extraction of membrane proteins into native lipid nanodiscs, enabling quantitative analysis of transient membrane-dependent protein complexes in sc-SiMPull experiments. Applying this approach to Cdc42/Par6 interactions, we discovered a remarkable modulation of the Cdc42/Par6 binding affinity between polarity establishment and maintenance, which are separated by only a few minutes of development. We also tested whether aPKC/Par6 can interact simultaneously with Cdc42 and Par3 as part of a 4-member complex, but found no evidence that such a complex exists *in vivo.* Together, our findings establish a novel approach for studying membrane protein interactions *in vivo* and reveal a new mode of polarity regulation in the *C. elegans* zygote.

## Results

### Cdc42 is necessary for Par6 recruitment to the plasma membrane during polarity maintenance but not polarity establishment

Motivated by recent biochemical work that has clarified the relationship between Cdc42, aPKC/Par6, and Par3 *in vitro* (Holly et al., 2020; Penkert et al., 2022; Vargas and Prehoda, 2023; Dickinson, 2023), we set out to measure the complexes these proteins form *in vivo.* We chose the *C. elegans* zygote as a model system for these experiments due to its stereotyped development, with distinct polarity establishment and maintenance phases in which polarity is thought to depend most strongly on Par3 or Cdc42, respectively (Gotta et al., 2001; Kay and Hunter, 2001; Cuenca et al., 2003; Beers and Kemphues, 2006; Aceto et al., 2006; Li et al., 2010a; b). We first wanted to confirm the general consensus in the field that Par3 and Cdc42 have distinct effects on Par6/aPKC localization during establishment compared to maintenance.

As expected, endogenously tagged Par6::mNeonGreen (mNG) and mNG::Par3 were localized at the anterior domain during polarity establishment (Figure 1A, left). During polarity maintenance, Par6 remained at the anterior cortex even though cortical levels of Par3 were strongly reduced, as previously reported (Figure 1A, right) (Dickinson et al., 2017). We then used RNA interference (RNAi) to deplete Cdc42, which reduced mNG::Cdc42 expression to undetectable levels (Figure 1B). Cdc42 depletion resulted in reduced cortical levels of Par6 during polarity maintenance, but not establishment, consistent with previous work (Figure 1A, 1D) (Gotta et al., 2001; Kay and Hunter, 2001; Aceto et al., 2006). Cdc42 depletion had no effect on Par3 levels at the anterior cortex during either establishment or maintenance (Figure 1A, 1C). Together, these results confirm that Cdc42 is required to stabilize Par6 at the anterior membrane during polarity maintenance but is dispensable for polarity establishment.

**Figure 1.**
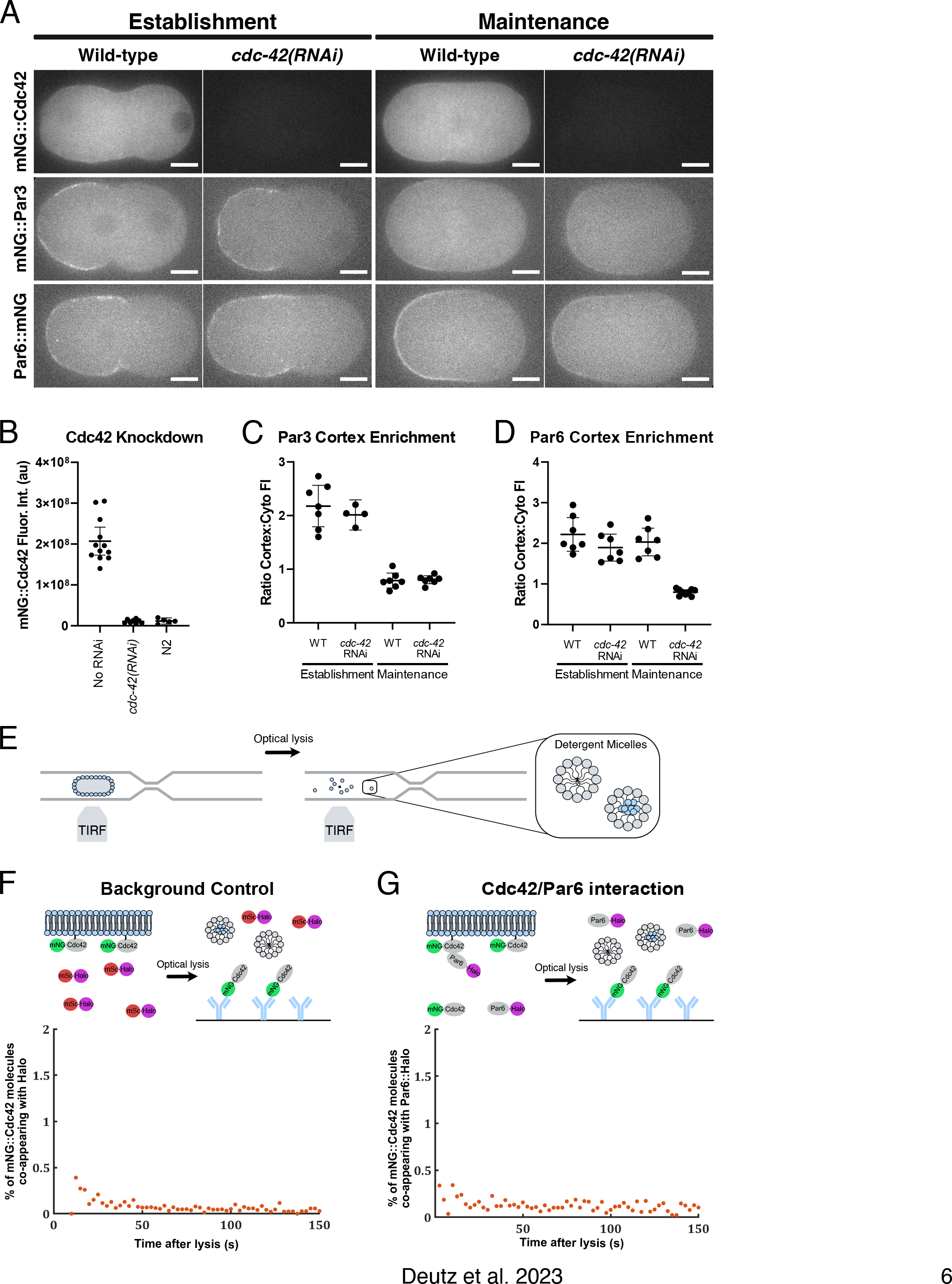
Cdc42 is necessary for Par6 recruitment to the plasma membrane during polarity maintenance, but their interaction is undetectable in detergent (A) Confocal images of *C. elegans* zygotes carrying endogenously tagged mNG::Cdc42 (top row), mNG::Par3 (middle row), or Par6::mNG (bottom row) during polarity establishment (left) or polarity maintenance (right), with or without *cdc-42(RNAi).* Scale bars represent 10 µm. (B) Quantification of whole-embryo mNG::Cdc42 fluorescence intensity in control (no RNAi) or *cdc-42(RNAi)* embryos, or in embryos with no mNG tag (N2). (C) Ratio of cortex to cytoplasm fluorescence intensity of mNG::Par3, from line scans taken during establishment (measured immediately after pseudocleavage furrow relaxation) or maintenance (measured after nuclear envelope breakdown) in either control (no RNAi) or *cdc-42(RNAi)* conditions. Each data point is a separate embryo. (D) Ratio of cortex to cytoplasm fluorescence for Par-6::mNG as described in (C). (E) Schematic of sc-SiMPull where a fluorescently tagged zygote is lysed in a microfluidic device and TIRF lasers are used to image bait (mNG::Cdc42) and prey (mSc::Halo or Par6::Halo) proteins as Cdc42 binds to a coverslip coated with anti-mNG nanobodies. (F) Fraction of mNG::Cdc42 molecules co-appearing with a non-interacting control protein (mSc::Halo) as a function of time, for experiments performed in detergent conditions (0.1% TX-100). N = 532,212 mNG::Cdc42 bait molecules counted from 8 embryos. (G) Fraction of mNG::Cdc42 molecules co-appearing with Par6::HaloTag as a function of time, for experiments performed in detergent conditions (1% TX-100). N=822,844 mNG::Cdc42 bait molecules counted from 8 embryos.

### The interaction between Cdc42 and Par6 is undetectable following cell lysis in detergent

Given the different genetic requirements for Cdc42 in establishment vs. maintenance, we wanted to interrogate whether the binding between Cdc42 and Par6 would also differ at different stages of polarity. To test this, we used a technique termed single-cell Single-Molecule Pull-down (sc-SiMPull) that can measure protein-protein interactions *ex vivo* after rapid single-cell lysis (Dickinson et al., 2017; Sarıkaya and Dickinson, 2021). Briefly, sc-SiMPull entails endogenously tagging proteins of interest using CRISPR-mediated knock-in and isolating a labeled cell (here, a *C. elegans* zygote) in a microfluidic channel (Figure 1E). The floor of the microfluidic device is a coverslip that is coated by antibodies, which recognize the fluorescent protein tag fused to one of the proteins of interest. Each cell is observed and imaged to record its developmental state. Then the cell is rapidly lysed using a pulsed infrared laser that generates a cavitation bubble, mechanically disrupting the cell without disrupting our ability to detect protein interactions (Sarıkaya and Dickinson, 2021). Immediately after lysis, TIRF microscopy is used to visualize proteins as they land on the coverslip and bind to antibodies.

When two proteins carrying different fluorescent tags land on the coverslip together, a simultaneous increase in fluorescence from both channels is observed, which is called co-appearance (Sarıkaya and Dickinson, 2021). We infer that co-appearing molecules are in complex with one another.

To detect the interaction between Cdc42 and Par6, we generated a strain with endogenously labeled mNG::Cdc42 and Par6::HaloTag. We labeled the HaloTag protein with the far-red dye JF_646_ (Grimm et al., 2015), lysed single staged zygotes, captured mNG::Cdc42 molecules, and counted the fraction that were in complex with Par6::Halo. As a negative control, we analyzed a strain carrying endogenously tagged mNG::Cdc42 along with a transgene expressing a fusion of mScarlet (mSc) covalently bound to HaloTag (mSc::HaloTag).

Unfortunately, despite the known interaction between Cdc42 and Par6 (Johansson et al., 2000; Joberty et al., 2000; Qiu et al., 2000; Garrard et al., 2003), we were not able to detect a specific association between these proteins under our standard assay conditions; the frequency of co-appearance between mNG::Cdc42 and Par6::HaloTag was similar to the background control (Figure 1F, 1G).

Our standard buffer for sc-SiMPull experiments, including those above, contains detergent (Triton X-100) to ensure solubilization of the plasma membrane following laser-induced cell lysis. Since Cdc42 is a lipid-anchored membrane protein, we suspected that the presence of detergent might disrupt its interactions with Par6/aPKC, or alternatively, that biologically active Cdc42/Par6 complexes reside in a membrane domain that is triton-insoluble. We tested several other detergents and varied the concentrations used, but were unable to find a condition that allowed the detection of Cdc42/Par6 complexes above background (Figure S1D, S1E). We, therefore, sought to develop an alternative method for studying membrane protein complexes in sc-SiMPull experiments.

### Nanodisc polymers rapidly solubilize membrane proteins from *C. elegans* embryos

The recent development of lipid nanodisc technology has allowed detergent-free solubilization of a wide variety of membrane proteins, which has been especially valuable for structural studies (Esmaili et al., 2021). Conventionally, lipid nanodiscs are formed by incubating purified proteins and lipids with amphipathic peptides (Bayburt et al., 2002; Bayburt and Sligar, 2010) or, more recently, polymer scaffolds (Knowles et al., 2009; Oluwole et al., 2017a; b) that encircle and stabilize a small lipid domain by forming hydrophobic interactions with the lipid tails. In one report, nanodisc-forming polymers were observed to clear the turbidity of a suspension of large unilamellar vesicles within a few seconds (Oluwole et al., 2017b), so we wondered whether these polymers would solubilize native cell membranes rapidly enough to be useful for sc-SiMPull experiments.

To test this idea, we lysed zygotes expressing a well-characterized plasma membrane marker, the pleckstrin homology (PH) domain from phospholipase-Cδ fused to mNG (Kachur et al., 2008; Heppert et al., 2016). We observed the lysis reactions using confocal rather than TIRF microscopy to allow visualization of the process of membrane dissolution. In control experiments without any solubilizing agent, the plasma membrane remained intact for more than 300 seconds after cell lysis (Figure 2A, first row). Addition of 1% Triton X-100 led to rapid dissolution of the membrane within a few seconds, as expected (Figure 2A, second row). Strikingly, the membrane was also solubilized within seconds by a nanodisc-forming diisobultylene/maleic acid (DIBMA) copolymer (Figure 2B, top row). We performed titration experiments with DIBMA and found that 1% w/v was required for rapid membrane dissolution (Figure 2B). Addition of 7.5mM CaCl_2_, which alters the size and stability of nanodiscs (Danielczak et al., 2019), resulted in membrane dissolution that was similarly fast (Figure 2B, bottom row). Rapid solubilization of the membrane was observed in the presence of another nanodisc-forming polymer, styrene/maleic acid 3:1 (SMA) (Bayburt and Sligar, 2010), but we did not pursue this reagent further because we found it contributed to unacceptable background fluorescence in TIRF experiments (SS, unpublished observations). Finally, we observed that the membrane pools of endogenously tagged Par polarity proteins were rapidly solubilized by DIBMA, including Cdc42 (Figure 2C), Par6 and aPKC (Figure S2).

**Figure 2.**
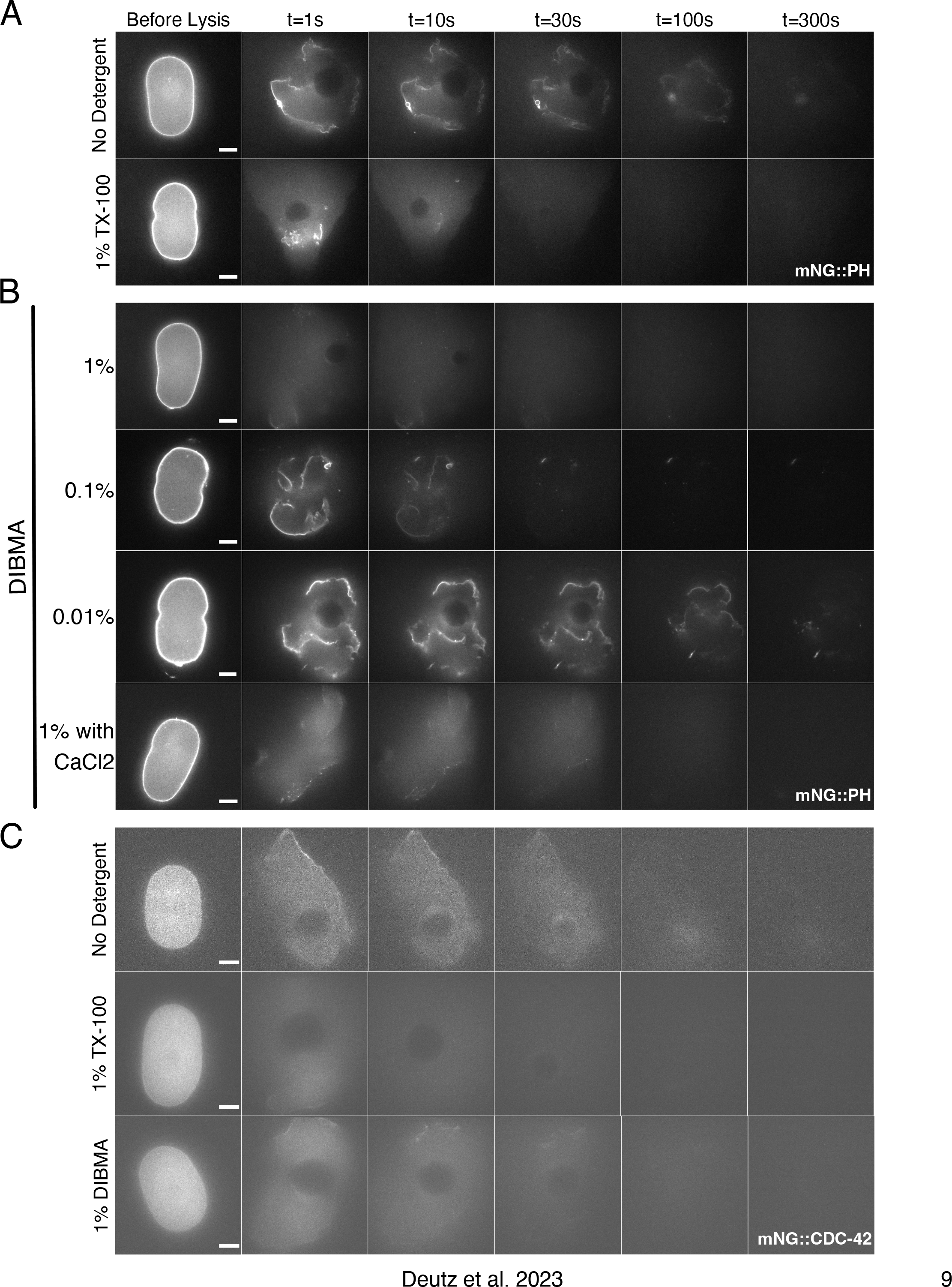
Nanodisc polymers rapidly solubilize membrane proteins from *C. elegans* embryos. All panels show confocal images of 1-cell embryos at the indicated times before and after rapid laser-induced cell lysis in the indicated lysis buffers. Images are representative of 3-10 embryos per condition. Scale bars represent 10 µm. (A,B) mNG::PH membrane marker. (C) Endogenously tagged mNG::Cdc42.

To verify that DIBMA was forming *bona fide* lipid nanodiscs in our experiments, we extracted the liquid from our microfluidic channels after laser lysis and visualized the lysate using negative-stain electron microscopy (Yi et al., 2019). In cells solubilized in detergent, we observed a heterogeneous population of protein particles, similar to what we have reported previously (Figure 3A) (Yi et al., 2019). In contrast, in the presence of 1% DIBMA buffer, we observed the formation of lipid nanodiscs (Figure 3B). Addition of divalent cations, which have been shown to modify nanodisc size *in vitro (Danielczak et al., 2019)*, led to the formation of larger nanodiscs under our conditions (Figure 3C–D). Native membrane nanodiscs formed in the absence or presence of CaCl_2_ had mean diameters of 33 nm and 42 nm, respectively, which is consistent with sizes reported for DIBMA nanodiscs formed from purified lipids *in vitro* (Figure 3D) (Oluwole et al., 2017a; b; Danielczak et al., 2019). No nanodiscs were present in DIBMA-containing buffer alone (in which no cell was lysed), confirming that lipid nanodisc formation is dependent on both the polymer and on cellular membrane lipids (Figure 3E). Together, these results indicate that DIBMA is capable of rapidly forming nanodiscs from native cellular membranes.

**Figure 3.**
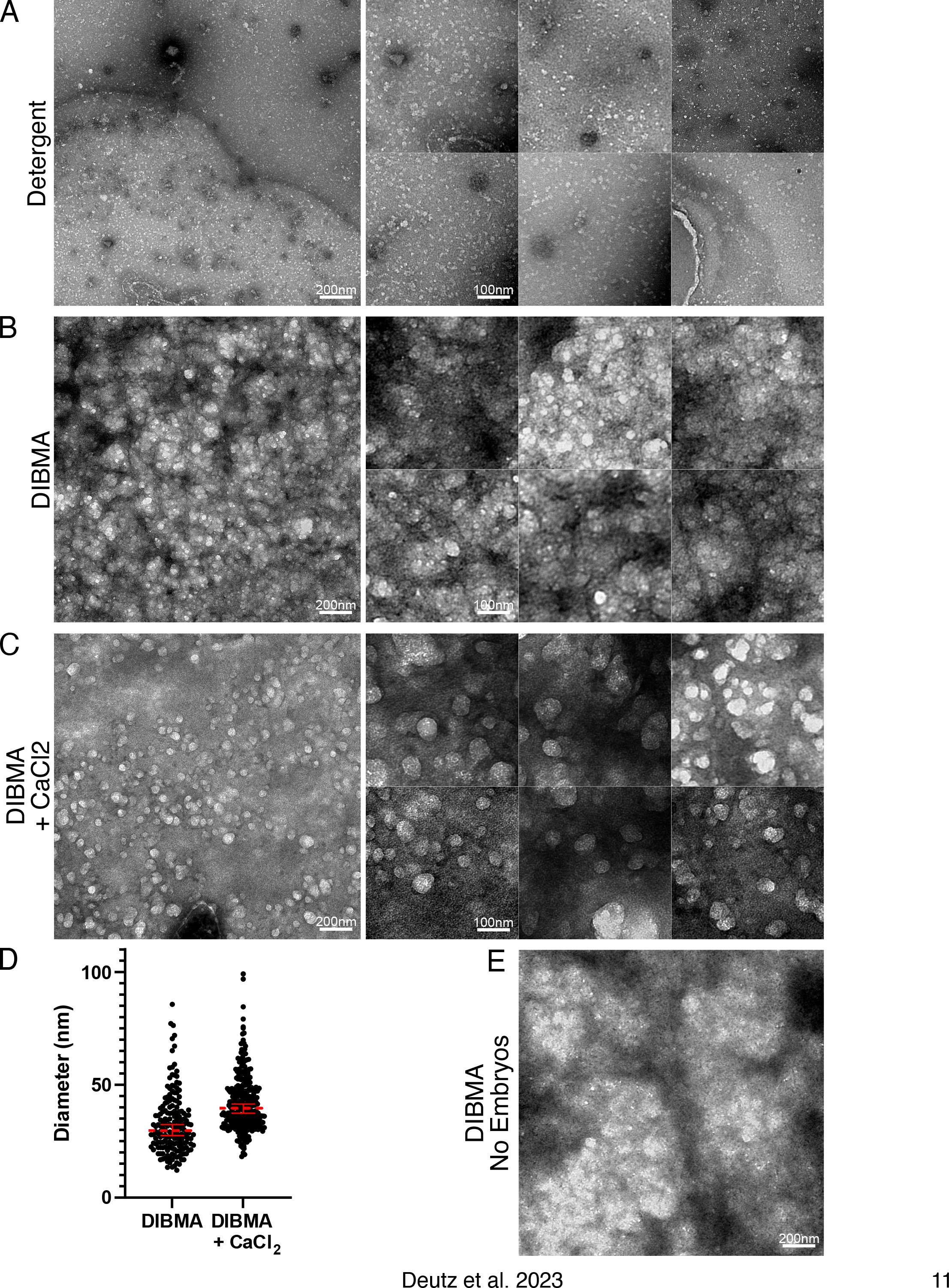
DIBMA polymers rapidly form lipid nanodiscs from *C. elegans* embryos. (A-C) Electron micrographs of cell lysate extracted from *C. elegans* embryos. Embryos were lysed in either (A) Detergent buffer, (B) 1% DIBMA, (C) 1% DIBMA with 7.5mM CaCl_2_, then extracted from the microfluidic channel, applied to an EM grid and negative-stained. (D) Quantification of average nanodisc diameter measured in either 1% DIBMA or 1% DIBMA with CaCl_2_. Dotted red line is the mean with error bars indicating the 95% confidence intervals. (E) Electron micrograph of DIBMA buffer on its own, with no embryos, after negative staining.

### Membrane extraction in lipid nanodiscs reveals temporal regulation of Cdc42 interaction with the Par complex

Returning to our original goal, we next asked whether native nanodisc formation would allow us to interrogate the Cdc42/Par6 interaction at different stages of development. We performed sc-SiMPull on staged embryos at either establishment or maintenance, using DIBMA-containing buffer. DIBMA did not increase the levels of background co-appearance measured using our negative control strain but did lead to robust detection of Cdc42/Par6 complexes (Figure 4A-C).

**Figure 4.**
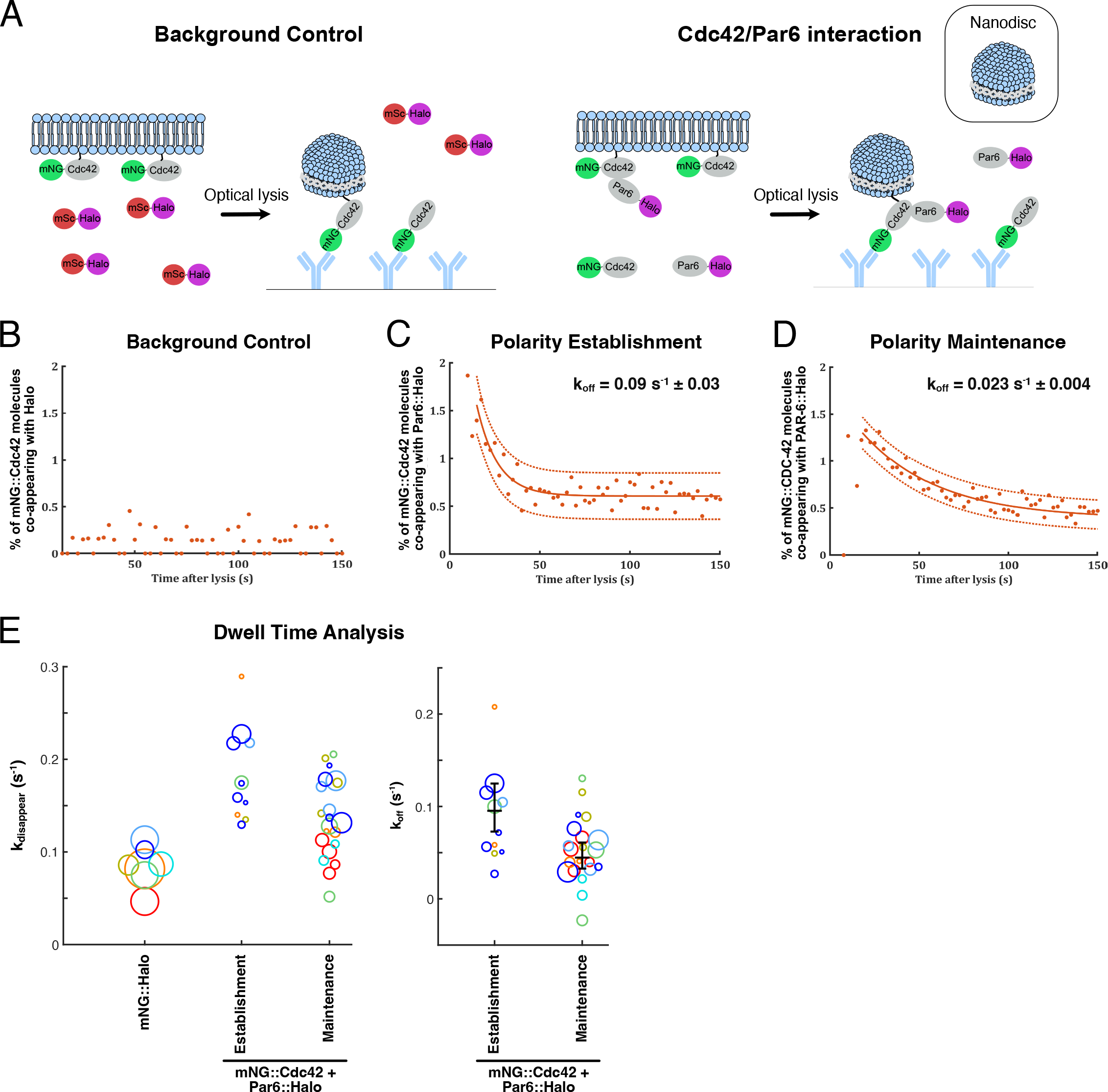
**The interaction of Cdc42 and Par6 is developmentally regulated.** (A) Schematic of the sc-SiMPull experiment with embryos being lysed in DIBMA buffer. (B) Fraction of mNG::Cdc42 molecules co-appearing with a non-interacting control protein (mSc::Halo) as a function of time, for experiments performed in 1% DIBMA conditions. N = 234,416 mNG::Cdc42 bait molecules counted from 6 embryos. (C-D) Fraction of mNG::Cdc42 molecules co-appearing with Par6::HaloTag as a function of time, for experiments performed in 1% DIBMA. Embryos were staged via brightfield microscopy immediately prior to lysis, and results from establishment- (C) and maintenance-phase (D) are shown separately. Curves were fit to single-exponential decay functions, resulting in the indicated estimates for k_off_. N = 2,753 mNG::Cdc42/Par6::HaloTag complexes from 17 embryos (Establishment); N = 7,746 mNG::Cdc42/Par6::HaloTag complexes from 34 embryos (Maintenance). (E) Calculation of k_off_ values for the Cdc42/Par6 interaction based on the distribution of single-molecule dwell times. Each data point represents one embryo, and the size of the circle represents the number of co-appearing signals analyzed from that embryo. Left panel: Measured rate constants for prey protein disappearance (k_disappear_) for mNG::Cdc42/Par6::HaloTag complexes and for the mNG::HaloTag control protein. Colors indicate experimental sessions (during which data was collected for 2 or more embryos). To account for day-to-day variation in laser power and system alignment, the mNG::HaloTag control data were used to calculate k_bleach_ for each experimental session, and that k_bleach_ was used to calculate k_off_ for Cdc42/Par6 experiments in the same session. Right panel: Calculated k_off_ values obtained by subtracting k_bleach_ from k_disappear_. Black lines show the weighted geometric mean and its 95% confidence interval. Embryos were staged via brightfield microscopy immediately prior to lysis. N = 1,516 mNG::Cdc42/Par6::HaloTag complexes from 11 embryos (Establishment); N = 4,043 mNG::Cdc42/Par6::HaloTag complexes from 21 embryos (Maintenance).

sc-SiMPull allows direct measurement of the stability of a protein-protein interaction. Native protein complexes dissociate after cell lysis due to dilution, and the speed with which this occurs can be analyzed to determine the dissociation rate constant k_off_ for an interaction (Sarıkaya and Dickinson, 2021). k_off_ can be measured in two different ways using the same data. First, we can monitor the fraction of co-appearing molecules as a function of time after lysis. Since the dissociation of protein complexes after dilution is a relaxation to equilibrium, the fraction of bait proteins co-appearing with prey declines to background levels following single exponential kinetics, which can be fit to extract the k_off_ (Sarıkaya and Dickinson, 2021). Remarkably, when we applied this analysis to Cdc42/Par6 complexes, we discovered that Cdc42/Par6 complexes captured from establishment-phase embryos dissociated nearly 4-fold faster than equivalent complexes extracted from maintenance-phase embryos (Figure 4C, D). We measured k_off_ = 0.09 ± 0.03 s^-1^ during polarity establishment and k_off_ = 0.023 ± 0.004 s^-1^ during polarity maintenance. If we assume a typical diffusion-limited k_on_ of ∼10^6^ M for this complex (Pollard, 2010), we estimate equilibrium binding affinities of 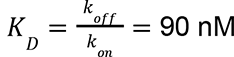 during polarity establishment and 23 nM during polarity maintenance. These values are in good agreement with the 50 nM binding affinity between Cdc42 and the Par6 CRIB-PDZ domain measured *in vitro* with purified proteins in one report (Garrard et al., 2003), though a more recent study reported a weaker affinity in the micromolar range (Vargas and Prehoda, 2023). It is striking that Cdc42 interacts more strongly with Par6 during maintenance phase, when Cdc42 is required for Par6 membrane recruitment, than during establishment phase when Cdc42 is dispensable.

An alternative, independent way to extract k_off_ from sc-SiMPull data is by analyzing the dwell times of single prey proteins (Sarıkaya and Dickinson, 2021). Dwell time is defined as the time between protein complex co-appearance and prey protein disappearance. From the distribution of dwell times, it is straightforward to calculate the disappearance rate constant k_disappear_ (Kinz-Thompson et al., 2016; Sarıkaya and Dickinson, 2021). Since prey protein molecules can disappear due to either unbinding or photobleaching, k_disappear_ = k_off_ + k_bleach_, where k_bleach_ is the photobleaching rate constant. Therefore, to estimate k_off_, we measure k_bleach_ from matched control experiments using a mNG::HaloTag fusion protein for which k_off_ = 0 (Figure 4E, left) (Sarıkaya and Dickinson, 2021). k_bleach_ is then subtracted from k_disappear_ to obtain k_off_ for the protein pair of interest. Applying this approach to our Cdc42/Par6 data, we determined k_off_ = 0.095 s^-1^ (95% confidence interval from 0.073 to 0.12 s^-1^) during polarity establishment and k_off_ = 0.045 s^-1^ (95% confidence interval from 0.033 to 0.061 s^-1^) during polarity maintenance, in good agreement with our results from fitting the relaxation curves for the entire population (Figure 4E, right; compare to Figure 4C, D).

It is important to emphasize that these two ways of calculating k_off_ from the data are independent and complementary. Relaxation curve fitting is a population analysis that considers the fraction of molecules found in a complex at different times following cell lysis, and ignores the lifetimes of individual complexes. The dwell time analysis is a single-molecule measurement that considers the bound lifetimes of individual complexes without regard to when (relative to lysis) those complexes were captured. Thus, the concordance between these two calculations increases our confidence that the faster k_off_ observed for Cdc42/Par6 complexes in establishment compared to maintenance phase is a real biological difference. Overall, we conclude that the *C. elegans* zygote actively regulates the stability of Cdc42/Par6 interactions, and these interactions are stronger during the stage of polarization when Cdc42 is found to have a stronger effect on Par6 localization.

### Par6 does not detectably interact with both Cdc42 and Par3 simultaneously

We next considered the relationship between Cdc42/Par6 complexes and the scaffold protein Par3, which plays a key role in transporting aPKC/Par6 to the anterior cortex during polarity establishment (Dickinson et al., 2017; Rodriguez et al., 2017; Chang and Dickinson, 2022; Illukkumbura et al., 2023). An early study found evidence that Cdc42, Par3, Par6 and aPKC could all be immunoprecipitated together in a 4-member complex (Joberty et al., 2000). However, the key experiment that supported a quaternary complex relied on overexpression of a GTP-locked Cdc42 mutant, and the same complex was not detected using endogenous Cdc42. More recent *in vitro* work has demonstrated that Par3 and Cdc42 can displace one another from aPKC/Par6 in solution (Vargas and Prehoda, 2023), indicating that aPKC/Par6 can bind to either Cdc42 or Par3, but not both simultaneously. Thus, it is unclear whether quaternary Cdc42/Par6/aPKC/Par3 complexes exist *in vivo*.

To test whether we could detect aPKC/Par6 complexes containing both Cdc42 and Par3, we constructed a triple-labeled strain carrying endogenously tagged mNG::Cdc42, Par6::HaloTag, and mSc::Par3. We then lysed single embryos, captured mNG::Cdc42, and looked for molecules that simultaneously co-appeared with both far-red (Par6::HaloTag) and red (mSc::Par3) signals (Figure 5A). We captured a total of 615,750 mNG::Cdc42 molecules from 20 embryos in these experiments, and found 41 molecules (an average of 2 per embryo) that appeared to contain all three signals. To determine whether these signals represent *bona fide* (albeit rare) Cdc42/Par6/Par3 complexes, we analyzed embryos carrying mNG::Cdc42 and Par6::HaloTag, but not mSc::Par3, under the same conditions. In this negative control experiment, mNG::Cdc42 co-appeared with both red and far-red signals at a similar frequency as in the experiment with all three proteins labeled (Figure 5B), suggesting that these signals are rare background artifacts, perhaps resulting from dust, fluorescence bleed-through or cellular autofluorescence. Thus, we could find no evidence that aPKC/Par6 forms complexes containing both Cdc42 and Par3 in *C. elegans* zygotes. We note that although this is a negative result, it is informative because we readily detected complexes of Cdc42/Par6 (Figure 4) and Par3/Par6 (Dickinson et al., 2017) in our assays. We conclude that aPKC/Par6 most likely forms separate complexes with Cdc42 and Par3 *in vivo*.

**Figure 5.**
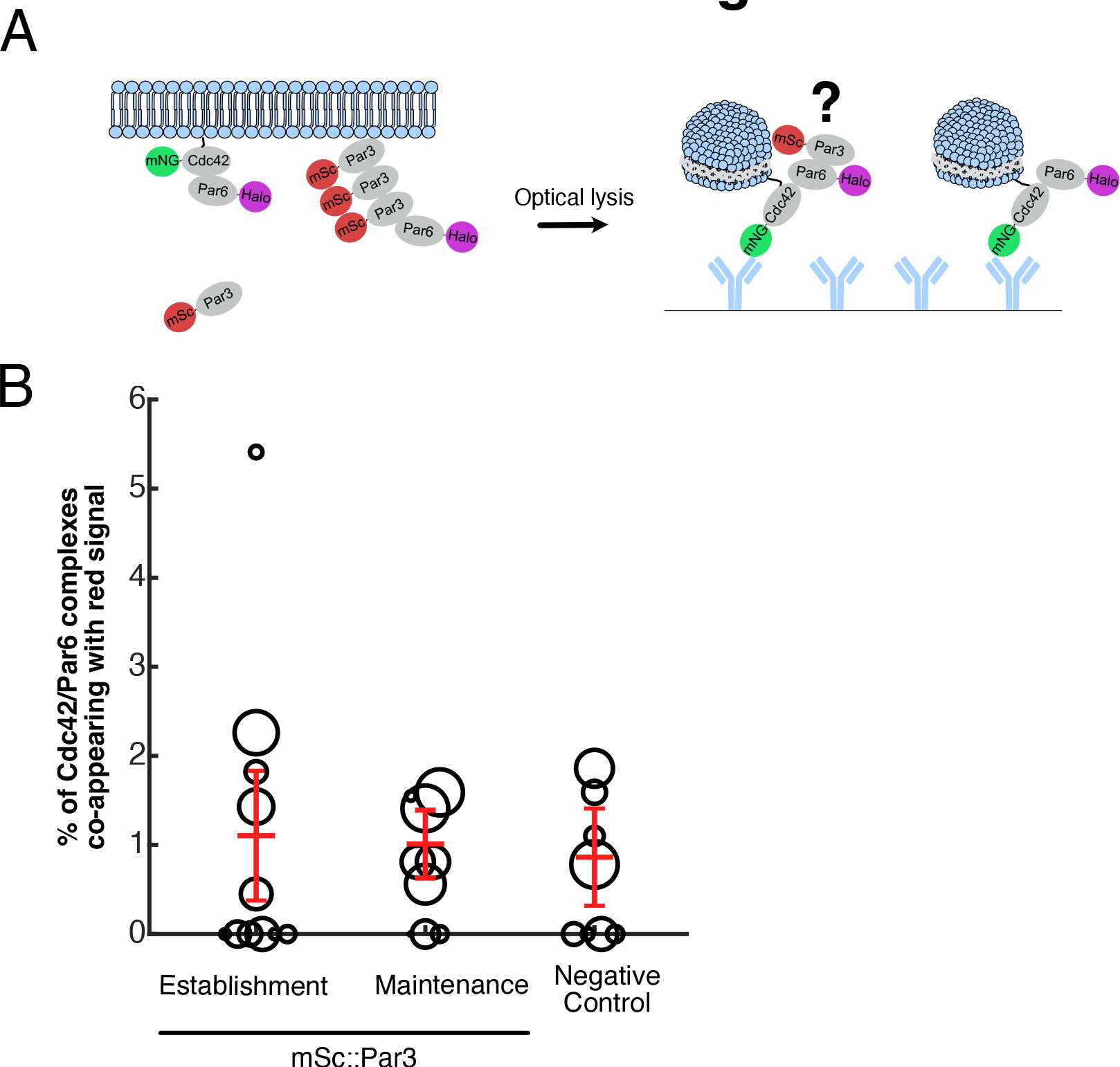
**Par6 does not interact simultaneously with Cdc42 and Par3.** (A) Schematic of the sc-SiMPull experiment: mNG::Cdc42/Par6::HaloTag complexes were captured and tested for co-appearance with mSc::Par3. (B) Fraction of mNG::Cdc42/Par6::HaloTag complexes co-appearing with mSc::Par3 in embryos lysed in 1% DIBMA conditions. Triple-labeled embryos (mNG::Cdc42; Par6::HaloTag; mSc::Par3) were staged via brightfield microscopy immediately prior to lysis, and results from establishment- and maintenance-phase embryos are plotted separately. Double-labeled negative control embryos (mNG::Cdc42; Par6::HaloTag) are mixed 1-cell stages. N = 1,720 mNG::Cdc42/Par6::HaloTag complexes from 11 embryos (Establishment); N = 2,115 mNG::Cdc42/Par6::HaloTag complexes from 9 embryos (Maintenance); N = 1,793 mNG::Cdc42/Par6::HaloTag complexes from 8 embryos (Negative Control).

## Discussion

Post-translational regulation of protein interactions is a hallmark of cellular signal transduction pathways and is essential for normal development. We have previously developed tools for studying the dynamic regulation of protein complexes at the single molecule level *in vivo*. In this study, we extend this approach to membrane proteins through the incorporation of amphipathic polymers that form native lipid nanodiscs. Using this approach, we demonstrate that the interaction between Cdc42 and Par6/aPKC is more stable during polarity maintenance compared to polarity establishment in the *C. elegans* zygote. This difference in complex stability is particularly striking considering that only a few minutes of development separate these stages. Our observations are consistent with genetic evidence indicating that Cdc42 does not play an essential role during polarity establishment but is required during polarity maintenance (Figure 1) (Kay and Hunter, 2001; Gotta et al., 2001; Aceto et al., 2006; Beers and Kemphues, 2006; Li et al., 2010b; a).

Cdc42, like other small GTPases, undergoes GTP-GDP cycling, and is considered active when in its GTP-bound form. Several studies have established that only active Cdc42-GTP binds to the aPKC/Par6 complex (Garrard et al., 2003; Vargas and Prehoda, 2023). These findings have led to the suggestion that local concentrations of GEFs and GAPs are the key factor that determines when and where Cdc42 recruits aPKC/Par6 to the plasma membrane.

Naively, this model seems to predict that there should be a greater number of Cdc42 molecules in complex with aPKC/Par6 during polarity maintenance compared to polarity establishment.

Surprisingly, we observed similar amounts of Cdc42-bound aPKC/Par6 complexes during both polarity establishment and polarity maintenance. Our measurements show that polarity entails regulation of the localization and affinity, rather than the total amount, of Cdc42/aPKC/Par6 complexes.

Although GEFs and GAPs undoubtedly regulate the distribution of Cdc42-GTP within the cell, regulation of GAP activity is unlikely to account for the developmental change in Cdc42/Par6 affinity that we observed. GEFs, GAPs, and effector proteins (such as Par6) bind to overlapping sites on small GTPases, such that a bound effector must dissociate from the GTPase before a GAP can catalyze GTP hydrolysis (Cherfils and Zeghouf, 2013). Thus, although GAPs could alter the (local) abundance of active Cdc42 in the cell, they would not be expected to catalyze the dissociation of effectors at the single-molecule level, as we observed for Par6 (Figure 4E). Instead, we hypothesize that Cdc42 may be post-translationally modified in a way that alters their binding affinity during either polarity establishment or maintenance. This hypothesis is supported by work in mammalian systems where it was found that phosphorylation states of Rho-family small GTPases, including Cdc42, can be an independent regulatory mechanism outside of GDP-GTP cycling (Forget et al., 2002). Work in cell culture has further suggested that phosphorylation of Rac1 and Cdc42 can shift the specificity of GTPase/effector coupling (Schwarz et al., 2012). Phosphorylation of Cdc42 is an attractive possible mechanism for tuning Cdc42/aPKC/Par6 affinity on the rapid timescales involved in *C. elegans* zygote polarization. Alternatively, it is possible that either Par6 or membrane lipids may be modified in a way that influences membrane recruitment of aPKC/Par6 by Cdc42. We intend to explore these ideas thoroughly in future work.

We also examined the relationship between Cdc42 and Par3, since these are the two key membrane-associated scaffolds that can recruit aPKC/Par6 to the cell cortex in the *C. elegans* zygote. Early work proposed the existence of a quaternary complex consisting of Cdc42, Par3, and aPKC/Par6 (Joberty et al., 2000), but more recent *in vitro* work suggests that aPKC/Par6 cannot bind to Par3 and to Cdc42 simultaneously (Vargas and Prehoda, 2023). We made thousands of observations of Cdc42/aPKC/Par6 complexes but found no evidence of a quaternary Cdc42/Par3/aPKC/Par6 complex. Of course, this does not rule out the existence of quaternary complexes in other cell types or under conditions we did not test. However, based on our results, we favor a model in which Par3 and Cdc42 recruit aPKC/Par6 to the plasma membrane in separate complexes, as previously proposed based on microscopy and genetic evidence (Hung and Kemphues, 1999; Beers and Kemphues, 2006; Rodriguez et al., 2017). We suspect that complexes containing both Par3 and Cdc42, if they exist at all, must be very transient and/or extremely rare.

The key technical innovation that allowed these biological insights was the use of maleic-acid copolymers to isolate membrane-associated complexes that are undetectable in detergent. The fact that Cdc42/aPKC/Par6 interactions are detectable in lipid nanodiscs but not in detergent is itself informative because it indicates that aPKC/Par6 requires an intact lipid bilayer in order to bind stably to Cdc42. This result is interesting in the context of two other recent findings. First, aPKC was found to bind directly to acidic phospholipids, indicating that membrane contact may influence aPKC’s conformation, localization, or activity (Dong et al.,2020; Jones et al., 2022). Second, the binding affinity of aPKC/Par6 for Cdc42 in solution, in the absence of any membranes, was found to be in the micromolar range (Vargas and Prehoda, 2023), which is approximately 100-fold weaker than what we estimate based on the k_off_ of the native complex. Based on these observations and our results, we propose that Cdc42/Par6 binding and aPKC/phospholipid interactions cooperate to recruit aPKC/Par6 to the plasma membrane. It will be important to test this hypothesis directly, using mutagenesis experiments and *in vitro* biochemistry.

Although we have not yet extensively explored complexes outside the PAR polarity system, we anticipate that the combination of sc-SiMPull and amphipathic nanodisc scaffolds will be broadly applicable to studying membrane protein interactions. We show that DIBMA is capable of solubilizing membranes within seconds following laser lysis, and this allowed us to detect transient Cdc42/Par6 complexes that have half-lives of less than 10 seconds (Figure 4C). Many membrane-associated signaling complexes have similar kinetics and thus might be amenable to this approach. As noted in previous work, sc-SiMPull can be broadly applied to any two proteins that can be fluorescently tagged, and the use of antibodies that recognize the fluorescent protein tags allows rapid application to different interactions without the need to re-optimize antibody binding conditions for each new protein pair. Notably, recent improvements in CRISPR-mediated fluorescent protein knock-in technology from our group (Shi et al., 2023) and others (Zhong et al., 2021) should facilitate the adaptation of this approach in mammalian systems.

As with any technique, there are limitations and outstanding technical challenges that need to be acknowledged. First, sc-SiMPull requires fluorescent tagging of proteins of interest. Endogenous tagging is preferred because it preserves native expression and ensures 100% labeling of the tagged proteins; however, sometimes fluorescent tagging can compromise gene function, and this needs to be assessed on a case-by-case basis for each newly generated fluorescent tag. Furthermore, we have observed cases in which steric hindrance between tags appears to prevent antibody binding, preventing us from capturing bait proteins. Lipid nanodisc formation could potentially compound this problem because a bulky nanodisc is more likely to sterically occlude the binding of an antibody to the tag. Secondly, some membrane proteins - particularly those that reside in unique lipid environments - might require different conditions for lysis or the use of alternative nanodisc-forming polymers. We have found it very useful to visualize the dissolution of membranes in real-time following laser lysis (as in Figure 2) in order to rapidly identify conditions that solubilize a protein of interest. Finally, our results to date suggest that it may be difficult to detect interactions that are much weaker than those we study here. A complex with micromolar affinity is expected to have a half-life of less than one second, which means that most of the cellular pool of protein complexes will dissociate before they diffuse to the antibody-coated surface and become detectable. In practice, this means that we will detect such complexes only if they are relatively abundant.

Despite these caveats, the extension of sc-SiMPull to membrane-bound proteins unlocks the study of a wide range of native signaling interactions that occur at the plasma membrane. In the example of Par polarity studied here, our experiments revealed native interactions between Cdc42 and aPKC/Par6, along with the regulation of these interactions on timescales of a few minutes *in vivo*. We emphasize that *in vitro* reconstitution remains a critical tool for basic studies of protein-protein interactions. However, as we show here, the advantage of sc-SiMPull compared to traditional measures of protein binding is our ability to reveal how protein interactions are modulated *in vivo* by developmental signals. Together, complementary *in vitro* reconstitution and *ex vivo* sc-SiMPull experiments promise to more clearly elucidate the mechanisms of cell signaling and behavior.

## Acknowledgements

We thank Luke Lavis for sharing JaneliaFluor dyes, and members of the Dickinson laboratory for helpful discussions and comments on the manuscript. Electron Microscopy was performed at the Center for Biomedical Research Support Microscopy and Imaging Facility at UT Austin (RRID:SCR_021756). This work was supported by an Undergraduate Research Fellowship from UT Austin (LND) and by NIH R01 GM138443 (DJD) and NSF MCB 2237451 (DJD). DJD is a CPRIT scholar supported by the Cancer Prevention and Research Institute of Texas (RR170054).

## Author Contributions

DJD conceived of the project, supervised the work and secured funding. LND and SS performed the experiments. LND and DJD analyzed the data using software written by SS and DJD. LND and DJD co-wrote the manuscript, and all authors discussed and contributed to the final version.

## Methods

### Materials, organisms and software

Antibodies, chemicals, strains and software are listed in supplementary table 1.

### *C. elegans* strain construction and maintenance

*C. elegans* were maintained on NGM medium and fed *E. coli* OP50 according to standard procedures. Fluorescent protein tags were inserted into the genome using CRISPR/Cas9-triggered homologous recombination, following published protocols (Dickinson et al., 2013, 2015).

### Whole-embryo imaging of *C. elegans* zygotes with RNA interference

RNA interference (RNAi) targeting *cdc-42* was performed by timed feeding. The RNAi feeding clone was retrieved from an RNAi feeding library (Kamath and Ahringer, 2003) and verified by Sanger sequencing before use. After being streaked out on an LB/Amp plate, individual cultures were grown in 4 mL of LB broth with ampicillin overnight. Cultures were subsequently concentrated into 1mL and 50-100uL of concentrated culture was spotted onto plates with 25 μg/mL Carbenicillin and 1 mM IPTG. Plates were left to dry for 4-24h at ambient temperature. Larval stage L4 worms were picked onto these plates, and embryos were dissected 18-24h afterward.

After being dissected from gravid adults, embryos were mounted with 22.8 µm beads (Whitehouse Scientific, Chester, UK) as spacers in egg buffer (5 mM HEPES (pH 7.4), 118 mM NaCl, 40 mM KCl, 3.4 mM MgCl_2_, 3.4 mM CaCl_2_). Images were acquired using a Nikon Ti2 inverted microscope (Nikon Instruments, Melville, NY) equipped with a 60x, 1.4 NA objective lens. Confocal images were taken with either an X-Light V3 spinning disk confocal head (Crest Optics, Rome, Italy) and a Prime95B sCMOS camera (Teledyne Photometrics, Tucson, AZ); or an iSIM super-resolution confocal scan head (Visitech, Sunderland, UK) and a Kinetix22 sCMOS camera (Teledyne Photometrics).

### Fabrication of microfluidic devices with antibodies

As described in previous work, microfluidic devices were fabricated using SU-8 photolithography (Stolpner and Dickinson, 2022). Briefly, a 10:1 mixture of PDMS:Curing agent was prepared, mixed, and poured onto molds. The devices were degassed for 2 minutes and afterward, transferred to a spin-coater set at 300 rpm for 30 seconds to ensure uniform ceiling height. Devices were left to bake in an 85c incubator for 20 minutes to solidify before being cut out to fit on coverslips and peeled off the mold. A 2 mm biopsy punch was used to punch out the inlet and outlet wells. 24x60mm glass coverslips were cleaned with compressed nitrogen gas and placed in a UV-Ozone cleaner for 20 minutes. PDMS devices were plasma treated and immediately placed in contact with the UV-Ozone cleaned coverslips to form a permanent bond.

A passivation solution was prepared by first dissolving 1% w/v of 3400 Da biotin-PEG-silane (Laysan Bio) in ethanol. 2 µL of this solution and 2 µL of water were added to 100 µL of liquid 600 Da mPEG-silane (Gelest inc.), resulting in a final concentration of 0.02% biotin-PEG-silane and 2% water. 2 µL of this mixture was placed in each inlet well and drawn into the channel under vacuum. 0.5 uL of PEG solution was subsequently placed in each outlet well and the devices were left to incubate at RT for 30 minutes. After incubation, devices were aspirated using a vacuum, rinsed 2x with water, and left overnight before use.

Immediately before an experiment, devices were functionalized with mNG nanobodies. A 0.2 mg/mL solution of Neutravidin in Tween buffer [10 mM Tris (pH 8.0), 50 mM NaCl, 0.1% Tween, and 0.1 mg/mL bovine serum albumin] was prepared and 1.5 µL was placed in each inlet wells and allowed to flow through the channel for ten minutes. After incubation, devices were rinsed 4x with Tween buffer, ensuring that the channel remained hydrated at all times. Subsequently, 1.5 µL of a 1 µM solution of biotinylated anti-mNG nanobodies in Tween buffer was placed in each inlet and allowed to flow through the channel for ten minutes. Devices were again rinsed with 4x with Tween buffer. After rinsing, 0.5 µL of Tween buffer was placed in each inlet and outlet, and devices were sealed with clear tape. Tape was cut with a scalpel and peeled off with forceps before each channel was used. Occasionally, devices were stored at 4°C overnight before use; this had no discernible effect on device performance.

### Labeling HaloTags with ligand dye

HaloTag ligand JF_646_ was dissolved in acetonitrile to a final concentration of 1 mM, dispensed into single-use 2 µL aliquots, dried and stored in the dark with desiccant at -20°C. The day before an experiment, aliquots were dissolved in 2 µL of DMSO to a concentration of 1 mM. 500 µL of an overnight culture of *E. coli* strain HB101 were spun down and resuspended in 100 µL of S-medium (150 mM NaCl, 1 g/L K2HPO4, 6 g/L KH2PO4, 5 mg/L cholesterol, 10 mM potassium citrate (pH 6.0), 3 mM CaCl2, 3 mM MgCl2, 65 mM EDTA, 25 mM FeSO4, 10 mM MnCl2, 10 mM ZnSO4, 1 mM CuSO4). Next, 1.5 µL of the dye mixture was added to obtain a final concentration of 15 µM HaloTag ligand dye. 30 µL of this mixture was added to one well of a 96-well plate and approximately 20 L4 larval stage worms were placed in the well. The plate was left to shake in a 25°C incubator overnight at 250 rpm.

### sc-SiMPull with TIRF microscopy

Gravid adults were dissected in egg buffer (5 mM HEPES (pH 7.4), 118 mM NaCl, 40 mM KCl, 3.4 mM MgCl2, 3.4 mM CaCl2) and rinsed 2x in either DIBMA buffer (10 mM Tris (pH 8.0), 150 mM NaCl, 1% DIBMA, and 0.1 mg/mL bovine serum albumin) or detergent buffer (10 mM Tris (pH 8.0), 50 mM NaCl, 0.1% Triton X-100, and 0.1 mg/mL bovine serum albumin). To rinse, embryos were moved with a mouth pipette into two 20 µL drops of lysis buffer. Separately, the microfluidic device channel was rinsed four times with lysis buffer to ensure no remaining tween was present. Embryos were subsequently transferred into the inlet well and either gently pushed into the channel with a clean 26G needle and/or pulled into the channel with vacuum.

Clear tape was used to seal both wells to prevent the channel from drying out and to prevent liquid flow. The microfluidic device with live zygote was transferred to the microscope and allowed to continue developing until optical lysis.

TIRF microscopy was performed using a custom-built microscope that utilized micromirror TIRF illumination (Friedman and Gelles, 2015) via the MadCity Labs RM21 platform (MadCity Labs, Madison, WI). The instrument was equipped with 488 nm, 561 nm and 638 nm excitation lasers; a 60x, 1.49 NA objective lens (Olympus, Tokyo, Japan); a home-built 4-color image splitter that allows simultaneous imaging of 4 wavelengths on a single camera chip; and a Photometrics PrimeBSI Express sCMOS camera (Teledyne Photometrics). After transfer to the microscope, embryos were located with transmitted light. A transmitted light image was captured to record the developmental stage of the embryo. The stage was moved ∼30 µm away from the embryo and TIRF focus was acquired. Using transmitted light, the stage was moved back to the embryo and the embryo was lysed with a single shot from a 1064 nm Q-switched Nd:YAG pulse laser (Teem Photonics, Meylan, France). After lysis, the stage was moved 60-120 µm from the point of lysis and TIRF images were acquired for 5,000-10,000 frames at 50 ms exposure. For more abundant proteins like mNG::Halo, the stage was moved 180 µm from the point of lysis to avoid an excessive density of molecules.

### Monitoring laser lysis with confocal microscopy

Embryos were prepared as described above, except that devices were not functionalized with antibodies. Images were acquired using a Nikon Ti2 inverted microscope equipped with a 60x, 1.4 NA objective; an iSIM super-resolution confocal scan head (Visitech); a Kinetix22 sCMOS camera (Teledyne Photometrics), and a optical lysis laser (Teem Photonics). Embryos were located with transmitted light. Time-lapse imaging was performed using a 488 nm laser to excite mNG fluorescence. Images were acquired every second for 300 seconds.

### Electron microscopy

Just before an experiment, several electron microscopy grids (Carbon Film on 200 mesh, Copper, PELCO® Pinpointer Grid) were plasma-treated. Wild-type embryos were prepared and lysed under the confocal microscope as described in *Monitoring laser lysis with* c*onfocal microscopy* with either No Detergent (10 mM Tris (pH 8.0), 50 mM NaCl), Detergent (10 mM Tris (pH 8.0), 50 mM NaCl, 1% TX-100), DIBMA buffer (10 mM Tris (pH 8.0), 50 mM NaCl, 1% DIBMA), or DIBMA with CaCl_2_ (10 mM Tris (pH 8.0), 150 mM NaCl, 1% DIBMA, and 7.5 mM CaCl_2_). After optical lysis, cell lysate was extracted from the microfluidic channel using a mouth pipette and a clean glass needle. Approximately 50 nL of cell lysate was transferred to a carbon-coated glow-discharged EM grid. The EM grid was subsequently negative stained with 35 µL of 2% (w/v) uranyl acetate solution and blotted to remove excess stain. The grid was imaged with a FEI Tecnai Transmission Electron Microscope (TEM) at 80kV.

### Quantification and Statistical Analysis

#### Fluorescence intensity measurements in intact embryos

To quantify Cdc42 depletion, whole-embryo fluorescence was measured by drawing a region of interest around the entire embryo and subtracting off-embryo background. To quantify enrichment of PAR proteins at the cortex, cortical and cytoplasmic fluorescence intensities were measured from line scans perpendicular to the cortex in the anterior part of the cell (Supplementary Figure 1B). Establishment embryos were measured right after furrow relaxation and Maintenance embryos after nuclear envelope breakdown. Measurements were made using FIJI, and scatterplots were prepared using Graphpad Prism.

#### Processing of dynamic sc-SiMPull data

SiMPull data analysis was performed using an open-source MATLAB package available at https://github.com/dickinson-lab/SiMPull-Analysis-Software/. The algorithms for spot detection, co-appearance detection, and kinetic analysis have been described in detail (Sarıkaya and Dickinson, 2021). Briefly, raw sc-SiMPull data consists of TIRF movies of single molecules binding to the antibody-functionalized coverslip. An image of sub-resolution fluorescent beads is collected on the same day as the experiment to allow automated registration of multicolor images produced by the image-splitting optics. The automated portion of the analysis performs image registration; generates a series of “difference images” by subtracting an average of each 50 frames from the average of the 50 frames that follows; segments the difference image to identify the locations of newly-appearing molecules; and then returns to the raw data and extracts an intensity-versus-time trace for each fluorescent channel at the location where a molecule appeared. Individual intensity traces are processed using a changepoint detection algorithm to determine the exact time when the fluorescence signal increases in each fluorescent channel. Simultaneous signal increases in fluorescence intensity for the bait and prey channels indicate co-appearance and are counted and tabulated by the automated portion of the analysis. We also implemented tools to automatically filter out known artifacts such as fluorescence bleed-through, excessive spot density and photoblinking (Sarıkaya and Dickinson, 2021).

Following the automated analysis, we manually inspected each imaging dataset to ensure that registration, spot detection and co-appearance detection were completed correctly, and we manually removed datasets or portions of datasets that contained out-of-focus images, excessive background fluorescence due to poor TIRF angle adjustment, macroscopic dust particles, and other artifacts.

Co-appearance versus time plots were generated by a plotting tool within the SiMPull analysis software package. Where appropriate, data were fit with a single exponential decay function to obtain k_off_ and its confidence interval (Sarıkaya and Dickinson, 2021). Counts of the number and fraction of co-appearing molecules, the total number of bait proteins counted, and k_disappear_ for each prey protein are outputs of the automated analysis and were tabulated in a spreadsheet where weighted means, k_off_ values and confidence intervals were calculated. We generally report weighted means (where the weight is proportional to the number of molecules considered in the analysis) for sc-SiMPull data, because each molecule is an independent observation, and weighting based on number of molecules ensures that each single-molecule observation contributes equally to the reported mean. The 95% confidence interval for the weighted mean is given by

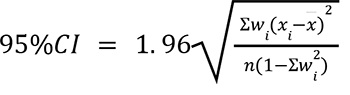

where *x*_*i*_ is the measured value from one single-embryo experiment, *x̄* is the weighted mean, *w*_*i*_ is the weight (defined as the number of molecules counted in experiment *i* divided by the total number of molecules counted in all experiments), *n* is the number of experiments performed, and the sums are over all experiments. Since we report geometric means for k_off_ values (which are log-normally distributed), we calculated the confidence intervals in Figure 4E using log-transformed data.

---

**Supplemental Figure 1.**
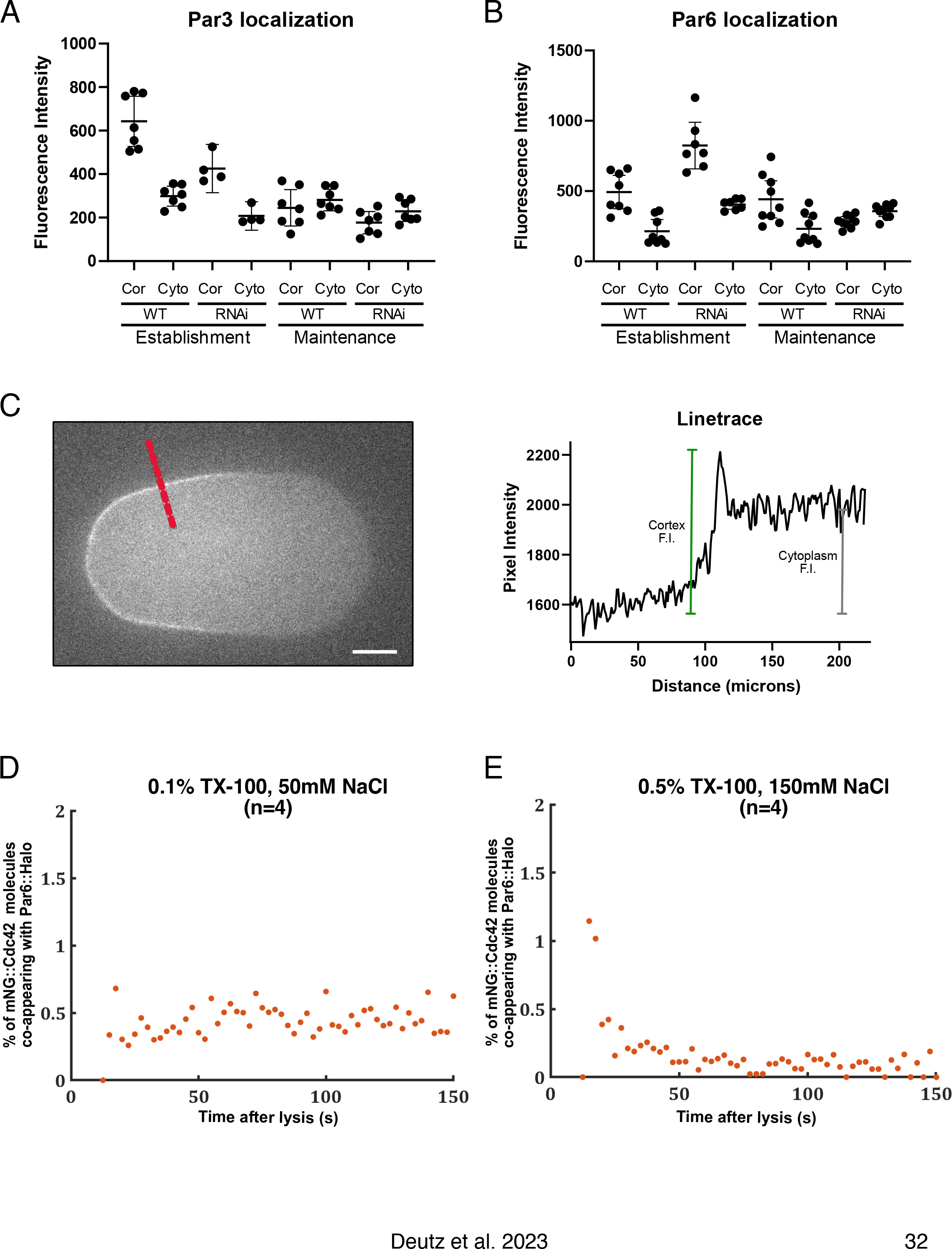
(A) Raw fluorescence values for mNG::Par3 as derived from linescans, separated by localization (cortex or cytoplasm), condition (wt or *cdc-42* (RNAi)), or developmental stage (polarity establishment or polarity maintenance) (B) Raw fluorescence values for Par6::mNG as described in (A) (C) Example of how fluorescence values for cortex and cytoplasm were measured from line scans. First, a line scan was drawn perpendicular to the embryo cortex (left) and a fluorescence by distance graph was generated (right). Cortex fluorescence was calculated as the maximum fluorescence at the peak at the edge of the embryo and the cytoplasmic fluorescence was calculated as an average of the embryo fluorescence. Off-embryo background was subtracted from both measurements. (D-E) Fraction of mNG::Cdc42 molecules co-appearing with Par6::HaloTag over time in a lysis buffer with less detergent (D, 0.1% TX-100) or a combination of more salt and intermediate detergent (E). N = 549,695 mNG::Cdc42 bait molecules counted from 4 embryos (Left); N=548,332 mNG::Cdc42 bait molecules counted from 4 embryos (Right).

**Supplemental Figure 2.**
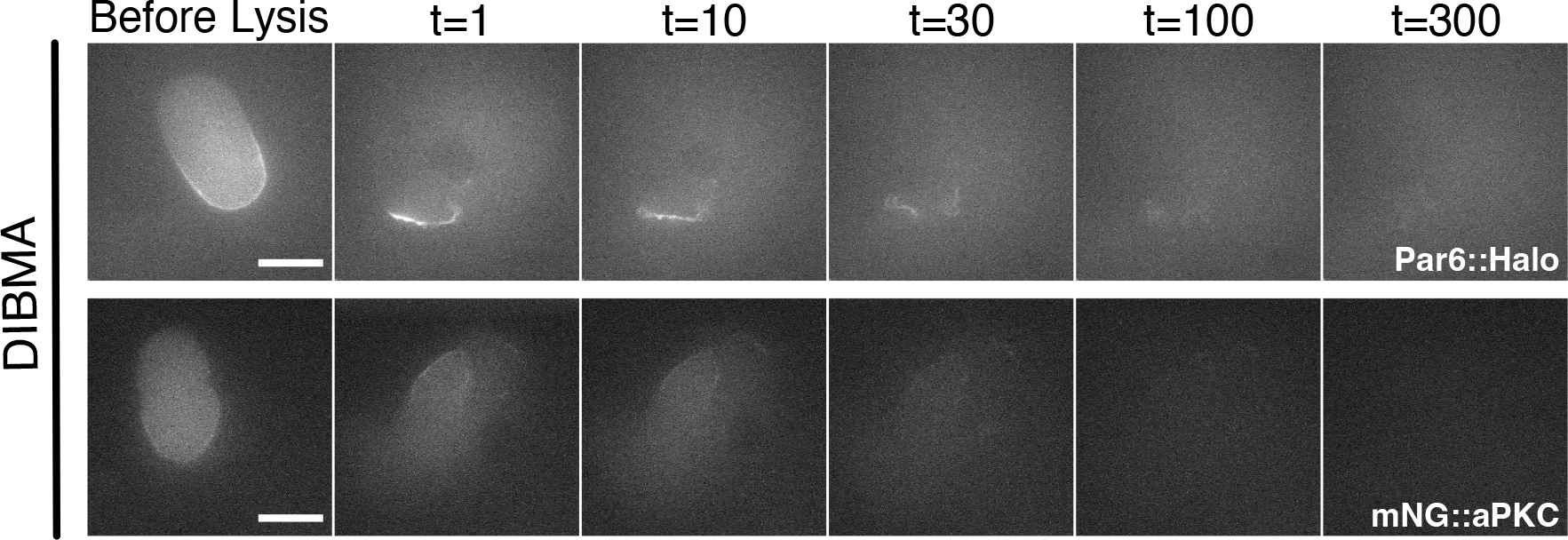
Confocal images of 1-cell embryos carrying the indicated endogenously tagged proteins at the indicated times before and after rapid laser-induced cell lysis in DIBMA-containing buffer.

